# Non-cell autonomous OTX2 in the piriform cortex regulates parvalbumin cell maturation states and olfactory-driven behavior

**DOI:** 10.1101/2024.02.04.578782

**Authors:** Rachel Gibel-Russo, Ariel A. Di Nardo

## Abstract

The timing of critical periods of juvenile brain plasticity is driven by the maturation of parvalbumin interneurons in the neocortex, a process regulated in part by non-cell autonomous activity of the OTX2 homeoprotein transcription factor. However, the involvement of critical periods in olfactory paleocortex maturation is unknown. Here, we find that the adult mouse piriform cortex parvalbumin interneurons display particularly low molecular maturation that increases in aged animals. Expression analysis of a large panel of genes reveals that an acute increase in piriform cortex OTX2 levels in young adult mice increases *Pvalb* expression as well as *Adamts9* expression, resulting in increased extracellular perineuronal net levels, while reducing OTX2 transfer decreases *Pvalb* expression and increases *Mmp9* expression, resulting in decreased perineuronal net levels. Reduction in OTX2 also stimulates odor-driven cFos activity in piriform cortex parvalbumin cells and disrupts olfactory-driven behavior. Our findings suggest plasticity in piriform cortex involves OTX2 activity on parvalbumin cells and lacks strictly defined critical periods.

## Introduction

Juvenile postnatal brain development involves critical periods of heightened plasticity that shape neural circuits in response to environmental stimuli and inherent biological constraints. These periods occur throughout the neocortex and are driven by the maturation of fast-spiking GABAergic interneurons expressing the Ca^2+^ buffering protein parvalbumin (PV) (Reh et al., 2020). Loss of plasticity after critical period closure is guided by a robust PV cell network resulting in consolidated circuitry maintained by molecular brakes, which include condensed extracellular matrix perineuronal nets (PNNs) surrounding PV cells (Carulli and Verhaagen, 2021; Fawcett et al., 2019; Testa et al., 2019). While PNNs are detected throughout the adult brain, some PV cells remain devoid of PNNs and show dynamic PV expression reflecting their level of activity for continued adult learning (Donato et al., 2013; Hensch, 2014). We sought to characterize the maturation and plasticity of PV cells in the piriform cortex (PCx), an associative region implicated in odor perception and odor-driven behaviors. While the rodent PCx shows sustained adult plasticity (Chapuis and Wilson, 2012; Wilson and Sullivan, 2011), in keeping with olfaction being a plastic sense requiring lifelong learning of odor associations, PCx PV cells can manifest PNNs (Ueno et al., 2019) and it is unclear whether critical period(s) occur in the PCx.

The PCx is a three-layered paleocortex with sensory-driven inputs from the olfactory bulb (OB) that lack defined topography (Bekkers and Suzuki, 2013; Giessel and Datta, 2014; Johnson et al., 2000; Klingler, 2017; Shepherd, 2011). It is divided into anterior (aPCx) and posterior (pPCx) regions that differ in connectivity and function (Neville and Haberly, 2004). The aPCx receives dense OB afferents and instructs odor identity, while the pPCx receives fewer OB afferents and participates in encoding odor qualities and associations. These circuits receive inputs from other olfactory areas, such as anterior olfactory nucleus and olfactory tubercle, as well as associative areas, such as medial prefrontal cortex (mPFC), hippocampus (HC), entorhinal cortex (EC), and basolateral amygdala (BLA), and also receive recurrent inputs from within the PCx. PV cells in the PCx are located in the deeper layers L2 and L3 and provide feedback inhibition to L1b, L2 and L3 pyramidal cells (Johenning et al., 2009; Suzuki and Bekkers, 2010). While their function in PCx circuitry is not completely understood, they have similar roles to neocortex feedforward PV cells for regulating spike activity, backpropagation, and induction of NMDA-dependent associative long-term potentiation (Canto-Bustos et al., 2022; Jiang et al., 2021; Johenning et al., 2009; Large et al., 2016).

In comparison to sensory regions of the neocortex, such as primary visual (V1) and auditory (A1) cortices, the PCx may not be expected to have well-defined critical period plasticity. Firstly, PCx cytoarchitecture is more similar to that of the HC, with feedback PV cells in the deeper layers of this three-layered structure (Shepherd, 2011). Secondly, the PCx lacks odor-driven topography and the sparsely activated unique ensembles of neurons drift in time independently of odor (Schoonover et al., 2021). Finally, the PCx receives the majority of its input from the OB, which undergoes continued circuitry remodeling through adult neurogenesis (Gheusi and Lledo, 2014). Thus, the PV cells within PCx are likely required to maintain a dynamic range in activity during adulthood, which appears in contradiction with the function of PNNs and their reported existence in PCx (Ueno et al., 2019). However, given that OB neurogenesis attenuates with aging, it may be that molecular brakes around PV cells may eventually develop and consolidate circuitry.

The timing of critical periods depends on the cortical region and sensory activity. For example, upon hearing onset, thalamocortical refinement occurs in A1 during a critical period from postnatal day (P) 12 to P15 in mice that can modify tonal response strength and topographic mapping (Takesian et al., 2018). The maturation of PV cells in V1, A1, and mPFC is regulated in part by the non-cell autonomous activity of the OTX2 homeoprotein transcription factor (Lee et al., 2017; Vincent et al., 2021). OTX2 activity promotes PV expression and critical period onset and also participates in critical period closure through induction of PNN expression (Beurdeley et al., 2012; Sugiyama et al., 2008). OTX2 can potentially reach all corners of the brain as it is secreted into the cerebrospinal fluid (CSF) by the choroid plexus and transferred specifically into cortical PV cells (Spatazza et al., 2013). OTX2 has been detected throughout the cortex but its accumulation has not been investigated in PCx.

We report that OTX2 is not only detected in the adult mouse PCx and but that its accumulation increases with age. By varying OTX2 levels in PV cells, in both aPCx and pPCx, we find that OTX2 does indeed impact PV cell activity and PNN accumulation by regulating the expression of PV and of PNN metabolic and catabolic enzymes. Reduction of OTX2 transfer into PCx PV cells enhances their activity and weakens olfaction-dependent memory. Thus, paleocortex feedback PV cells can be controlled by OTX2 similar to neocortex feedforward PV cells, suggesting that age-dependent plasticity states exist in the PCx without strictly defined critical periods.

## Results

### PV cell maturation kinetics depends on brain structure and cell subtype

In order to facilitate quantification of PV intensity, we initially turned to the oft-used *PV::Cre* × *Rosa-tdTomato* (*PV-tdTom* hereafter) mouse line for genetic identification of PV cells independently of PV expression level (Donato et al., 2013; Tang et al., 2014). As PV is expressed in various regions throughout the brain, including V1, HC, BLA, mPFC and the PCx (Celio, 1990), we first sought to compare the distribution and maturation level of PV cells across these regions in adult P90 mice (Figure 1A-C). In *PV-tdTom* mice, the tdTomato red fluorescent protein (TOM) localizes throughout the cell and its expression level is not correlated with that of PV. We found a higher density of PV+ cells in V1, and a higher density of TOM+ cells in V1 and mPFC compared to the other regions (Figure 1B). Within the HC, we detect more PV+ and TOM+ cells in CA1-3 compared to the dentate gyrus (DG). The density of PV cells is clearly dependent on brain region, in agreement with previous studies (Bjerke et al., 2021; Kim et al., 2017). To evaluate the maturation level of PV cells in each region, we compared the number of TOM+ cells stained for PV+ to the total number of TOM+ cells (Figure 1C), since this allows to take into account “immature” PV cells not expressing PV, i.e. PV(-)TOM+ cells. We found PV cells in V1, BLA and CA1-3 to be significantly more “mature” than in DG, mPFC and PCx. To further evaluate PV cell maturation, we also performed WFA staining which detects the PNNs around “mature” PV cells (Figure 1C). Although this technique relies on detecting PV expression and will thus miss fully “immature” PV cells, such as the PV(-)TOM+ cells, the pattern of PV cell maturation across regions was found to be similar to the pattern observed with the TOM staining method. Thus, we consider the PV+TOM+ and WFA+PV+ methods equally robust for use as a proxy to quantify PV cell maturation levels. Given that only approximately 30% of PV cells in the PCx are “mature” (Figure 1C), circuit inhibition driven by PV cells is likely less strong in the PCx compared to V1 and CA1-3 of P90 mice.

**Figure 1.**
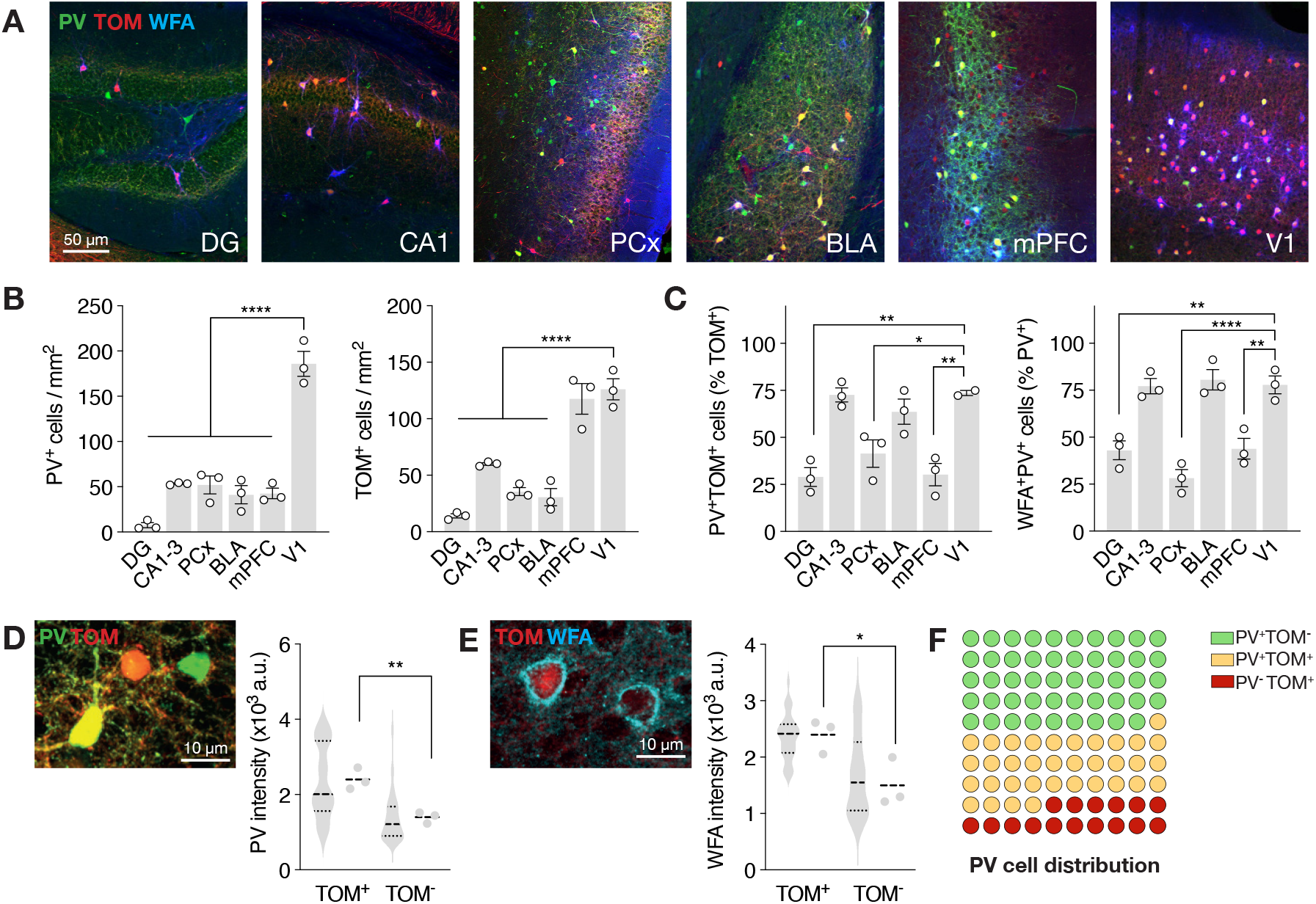
PV cell maturation kinetics depends on brain region and cell subtype in the PCx. **(A)** Representative images of PV, TOM and WFA staining in the Hc (DG and CA1), PCx, BLA, mPFC, and V1 at P90 in *PV-tdTom* mice. **(B)** Number of PV and TOM positive (+) cells in the Hc (DG and CA1-3), PCx, BLA, mPFC, and V1 at P90 in *PV-tdTom* mice. **(C)** Percentages of TOM+ cells stained for PV and PV+ cells stained for WFA in the Hc (DG and CA1-3), PCx, BLA, mPFC, and V1 at P90 in *PV-tdTom* mice. **(D)** Representative image and quantification of PV staining intensity in TOM negative (TOM-) and TOM+ cells in PCx. Violin plots show values for individual cells, while adjacent data points represent the average cell intensity per mouse. **(E)** Representative image and quantification of WFA staining intensity in TOM- and TOM+ cells in PCx. **(F)** Distribution of TOM+ and TOM- cells in the identified PV cell population. All values ± SEM; n = 3 mice per group; one-way ANOVA, post hoc Tukey’s or Dunnett’s test in (B, C); *t*-test in (D, E); \**P* < 0.05, \*\**P* < 0.01, \*\*\**P* < 0.001.

It is known that not all PV cells express TOM in *PV-tdTom* mice (Madisen et al., 2010), that *PV::Cre* mice provide low efficiency labeling in some brain regions (Nigro et al., 2021), and that epigenetic silencing of transgenes may occur at the ROSA26 locus in adult mice (Gödecke et al., 2017). However, a recent study suggests there may be a high amount of overlap between TOM and PV expression in the aPCx of adult *PV-tdTom* mice, although data was reported for one animal only (Large et al., 2016). To determine the level of heterogeneity in the PCx PV cell population, we calculated the level of PV staining intensity in both PV+TOM+ and PV+TOM(-) cells (Figure 1D), knowing that PV intensity and TOM staining are expected to be independent as they are reside on distinct gene loci. We observed a bias of PV staining intensity toward the TOM+ population, suggesting that the most mature cells express TOM. We also observed a similar bias for the intensity of WFA (Figure 1E), suggesting that the TOM+ cell population is indeed a distinct cellular subpopulation of PV cells. Furthermore, only ~50% of the PV cells express TOM in the PCx of P90 mice (Figure 1F), suggesting developmental divergence and/or adult ROSA26 silencing (Gödecke et al., 2017; Madisen et al., 2010). Given that the purpose of this study was to evaluate PV cell maturation in the PCx, a region for which this has not been previously evaluated, we chose not to use *PV-tdTom* mice for subsequent analyses in order to avoid potential bias in cell subtype. Nevertheless, the *PV-tdTom* mouse model provides the advantage of detecting PV cells with very low PV expression levels, and in the adult PCx ~30% of TOM+ cells have undetectable levels of PV staining (PV(-)TOM+ group in Figure 1F). By ignoring this cell population, we may be missing information regarding the immature low-activity PV cells. However, this population only represents ~16% of detected PV cells in the PCx (Figure 1F), and we determined that the alternative WFA+PV+ method remains pertinent as a proxy for detecting mature PV cells (Figure 1C). Thus, the quantification of PV staining levels in the PCx provides a fairly accurate picture of PV cell maturation state despite not taking into account PV cells with undetectable levels of PV expression.

### Piriform cortex matures late and depends on non-cell-autonomous OTX2

Compared to V1 in which PNN formation is highly enriched after CP closure (Lee et al., 2017; Pizzorusso et al., 2002), we observed significantly lower PNN enrichment in the PCx of adult mice (Figure 1C). Given that PNNs around PV cells are a hallmark of consolidation and reduced plasticity, we hypothesized that this associative 3-layered cortex remains plastic at P90. To more finely characterize PV cells, we separated analysis between the aPCx and posterior PCx (pPCx) subregions, given their different cytoarchitecture and connectivity (Neville and Haberly, 2004), and focused on cortical layers 2 and 3 given the absence of PV cells in layer 1 (Ekstrand et al., 2001; Suzuki and Bekkers, 2010). We also analyzed PV cells in P400 mice to determine whether the PV expression kinetics beyond P90 is either stable or continues to mature. For both ages, we observed no differences between the aPCx and pPCx, but found a significant increase in the number of PV cells in both aPCx and pPCx at P400 (Figure 2A). Throughout the brain, PV cells are a heterogeneous population with varying degrees of PV expression showing different stages of maturity. Previous studies in the HC have shown that low or high PV states are associated with increased GABAergic or excitatory synaptic inputs to the PV cells themselves, respectively (Donato et al., 2013). The lower PV states correspond to higher plastic states while the higher PV states correspond consolidated non-plastic states. We defined classes based on the level of PV staining intensity and found that the distribution of PV cell classes is similar between aPCx and pPCx and depends on age (Figure 2B). In younger animals, the PV cell group with the lowest PV intensity (PV-low) represents almost 50% of the total number of PV cells, whereas in older animals, PV-low is the least represented class (Figure 2B).

**Figure 2.**
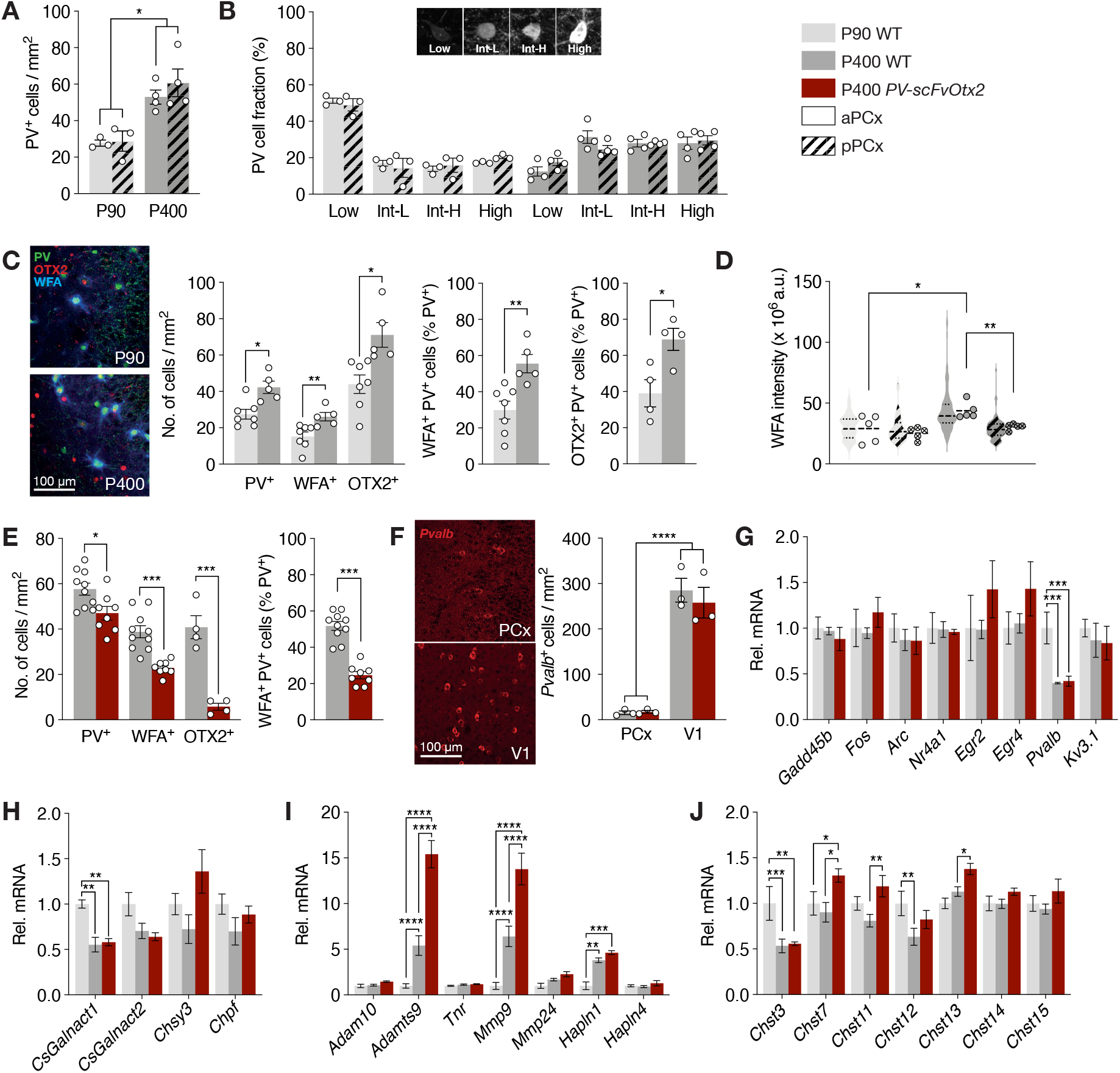
The piriform cortex matures late and depends on non-cell-autonomous OTX2. **(A)** The number of PV+ cells in aPCx and pPCx at P90 and P400 (n = 3-4 mice per group). **(B)** The distribution of the level of PV staining intensity in aPCx and pPCx at P90 and P400. Inset shows example images of PV staining intensity for each category (from left to right: low-PV staining, intermediate-low-PV staining, intermediate-high-PV staining, high-PV staining; n = 3-4 mice per group). **(C)** Representative images and quantification of the number of PV+, WFA+, OTX2+ cells and percentage of PV+ cells stained with either WFA or OTX2 (n = 4-7 mice per group). **(D)** The intensity of WFA staining in PV+ cells (n = 5 mice per group). Violin plots show values for individual cells, while adjacent data points represent the average cell intensity per mouse. **(E)** The number of PV+, WFA+, and OTX2+ cells the number of PV+ cells stained for WFA in P400 WT and *PV-scFvOtx2* mice (n = 4-10 mice per group). **(F)** Representative images and quantification of the number of cells expressing *Pvalb* mRNA in PCx and V1 layer IV of WT and *PV-scFvOtx2* P400 mice (n = 3 mice per group). **(G)** Quantitative PCR analysis of plasticity-related genes and *Pvalb* (n = 4-5 mice per group). **(H)** Quantitative PCR analysis of CSPG synthesis genes (n = 4-5 mice per group). **(I)** Quantitative PCR analysis of ECM components (n = 4-5 mice per group). **(J)** Quantitative PCR analysis of *Chst* genes (n = 4-5 mice per group). All values ± SEM; one-way ANOVA, post hoc Tukey’s (A, D, F-J); two-way ANOVA in (B); *t*-test in (C, E); \**P* < 0.05, \*\**P* < 0.01, \*\*\**P* < 0.001.

Since OTX2 is secreted from the choroid plexus into the CSF, interacts with PNNs, and is internalized by PV cells in several cortices (Lee et al., 2017; Spatazza et al., 2013; Vincent et al., 2021), we hypothesized it could be involved in PCx maturation. OTX2 accumulation was indeed detected in both aPCx and pPCx, and, similar to PV staining, both OTX2+ and WFA+ cell numbers increased with age (Figure 2C). Furthermore, the percentages of PV+ cells co-stained with WFA and the percentage of OTX2+ cells co-stained with PV also increased in P400 mice (Figure 2C). Taken together, these results show that PV expression, PNN assembly, and OTX2 accumulation is age-dependent, and that the PCx undergoes late maturation compared to other cortices. However, while the kinetics of the distribution of PV staining intensity do not differ between aPCx and pPCx, we observed a significant increase in WFA staining intensity at P400 compared to P90 only in the aPCx (Figure 2D), suggesting that these sub-regions mature differently.

To investigate the role of OTX2 in critical periods, we previously generated the *PV::Cre;scFvOtx2*^*tg/o*^ knock-in mouse line (*PV-scFvOtx2* hereafter) in which a neutralizing single-chain variable fragment (scFv) against OTX2 is secreted from PV cells and traps OTX2 in the extracellular milieu (Bernard et al., 2016). Blocking extracellular OTX2 reduces its accumulation around PV cells and subsequent internalization, which was shown to delay critical period onset in V1 (Bernard et al., 2016). To follow PV cell maturation in the PCx, P400 sections were stained for OTX2, PV, and WFA (Figure 2E). The expression of scFv-OTX2 reduces the number of PV+ cells by ~30%, the number of WFA+ cells by ~50%, and the number of OTX2+ cells by ~75%. Furthermore, the number of PV+ cells that have PNNs (WFA+PV+) is significantly reduced compared to WT (Figure 2E). To verify whether the reduction in the number of PV-stained cells was not due to a loss of PV cells, we performed RNAscope in situ hybridization (ISH) to more sensitively identify cells expressing *Pvalb* RNA, comparing V1 to PCx in both WT and *PV-scFvOtx2* mice (Figure 2F). We found no difference in the number of cells expressing *Pvalb* RNA, suggesting that the decrease in PV cell number in P400 *PV-scFvOtx2* mice is due to a relative increase in the PV-low population that cannot be detected by PV staining. Thus, blocking extracellular OTX2 delays the maturation of the PCx suggesting that OTX2 could act as a regulator of PCx plasticity as it does in V1, A1, and mPFC (Lee et al., 2017; Vincent et al., 2021).

To determine whether the changes in PV and WFA staining observed at P400 in both WT and *PV-scFvOtx2* animals are due to altered expression of plasticity-related genes and genes related to PNN components, we quantified mRNA levels from micro-dissected PCx (aPCx and pPCx combined). The expression of plasticity-related genes did not differ between P90 and P400 WT mice, nor between P400 WT and *PV-scFvOtx2* mice (Figure 2G). However, *Pvalb* expression decreased by more than 50% at P400, in both WT and *PV-scFvOtx2* mice, suggesting long-term scFvOTX2 expression does not impact *Pvalb* expression (Figure 2G). However, as PV staining reveals reduced protein levels in the *PV-scFvOtx2* mice, *Pvalb* mRNA and PV protein levels are clearly not correlated, and regulation is therefore likely at the translation level. Such regulation is also evident in WT mice, given that the number of PV-stained cells increase between P90 and P400 WT mice while *Pvalb* mRNA levels decrease.

### OTX2 affects the expression of PNN components in the piriform cortex

Given that both age and OTX2 can affect PNN levels, we sought to determine whether various PNN-related enzymes and components are differentially expressed at P400 in either WT or *PV-scFvOtx2* mice. PNNs are dynamic ECM structures composed of hyaluronan, chondroitin sulfate proteoglycans (CSPGs), and scaffolding proteins that typically condense around PV cells (Fawcett et al., 2019). Of the enzymes involved in CSPG biosynthesis that we examined, only *CsGalnact1* was found to be differentially regulated at P400, in both WT and *PV-scFvOtx2* mice, pointing to no effect of scFvOTX2 on their expression (Figure 2H). Regarding PNN structure and turnover, the ADAM and MMP families of extracellular proteases are often involved in ECM remodeling (Huntley, 2012; Kelwick et al., 2015), while tenascin-R (TNR) and the HAPLN family are involved in PNN stabilization (Deepa et al., 2006). Among the various genes we analyzed, *Adamts9, Mmp9*, and *Hapln1* are the only ones with significant increases in gene expression between P90 at P400 WT mice (Figure 2I), suggesting that the PCx PNN is reorganized in aged mice. The proteases and HAPLN1 should have opposing effects on PNN stability, but the net effect is an increase in PNNs with age (Figure 2C). On the other hand, *PV-scFvOtx2* mice have increased *Adamts9* and *Mmp9* expression compared to P400 WT mice, with no change in *Hapln1*, which could explain the observed decrease in PNNs in these mice (Figure 2E).

Because OTX2 preferentially recognizes di-sulfated CSPGs and participates in a feedback loop to increase PNN expression (Di Nardo et al., 2018), we also quantified the expression of the chondroitin sulfate transferase (Chst) family (Figure 2J). These enzymes add sulfates to *N*-acetylgalactosamine (GalNac) of chondroitin sulfate (CS) (Mikami and Kitagawa, 2013), leading to several CSPG subtypes for which the pattern of sulfation defines their charge structure, binding affinities, and biological functions (Kitagawa et al., 1997; Sugahara and Mikami, 2007; Testa et al., 2019). CHST3 and CHST7 sulfate at GalNac position 6 (6S), while CHST11, CHST13, and CHST14 sulfate at GalNac position 4 (4S), with subsequent sulfation on position 6 by CHST15 resulting in 4S/6S di-sulfated CS. The level of 6S has been shown to decrease with age, while the 4S/6S ratio increases with cortical maturation (Miyata et al., 2012). Only *Chst3* has altered expression and decreases in P400 WT mice (Figure 2J), which is consistent with decreasing 6S levels in neocortex with age (Miyata et al., 2012). A similar decrease is also observed in *PV-scFvOtx2* mice, suggesting scFvOTX2 does not impact CHST3 levels. However, these mice also experience increased expression of *Chst7, Chst11*, and *Chst13*, which provide mono-sulfated CSPGs (CS type-A and type-C) and the precursors for di-sulfated CSPGs (CS type-D and type-E). We observe no change in *Chst15*, which provides di-sulfated CSPG (type-E) that bind OTX2 (Beurdeley et al., 2012; Miyata et al., 2012). Nevertheless, taken together, these results suggests that the decreased OTX2 uptake by PV cells in *PV-scFvOtx2* mice is very likely affecting sulfation patterns in the ECM of the PCx.

### The piriform cortex undergoes remodeling after acute reduction of OTX2 transfer

A caveat of the *PV-scFvOtx2* mouse model is that it results in brain-wide disruption of OTX2 transfer into PV cells, potentially affecting plasticity across the brain at different times, depending on the region. Given the associative function of the PCx, the delay in PV cell maturation observed with the *PV-scFvOtx2* mouse could be due to additive effects from changes in connected regions, such as EC, BLA, and mPFC. To ensure that altered maturation is due in part to intrinsic changes, we performed acute OTX2 loss-of-function locally in PCx. Double-floxed inverse open (DIO) reading frame AAV8-EF1a-DIO-GFP-2A-scFvOtx2 virus was injected in PCx of *PV::Cre* mice for conditional PV-cell-specific expression, with GFP expressing-neurons detected in both aPCx (+1.03 ± 0.28 from bregma) and pPCx (−1.84 ± 0.46 from bregma), 3 weeks after P90 injection (Figure 3A). As with the *PV-scFvOtx2* mice at P400, we observed a decrease in PV+ cells, along with a ~50% decrease of WFA+ cells accompanied by a dramatic reduction of OTX2+ cells (Figure 3B). PV cell maturation level was reversed in both aPCx and pPCx, as assessed by the decrease of WFA staining around PV cells (Figure 3C). The distribution in PV staining intensity was shifted towards the low-intensity population, with a more marked loss of higher-intensity populations in the pPCx (Figure 3D). Furthermore, not only does the number of WFA+ cells decrease after scFvOTX2 expression (Figure 3B), but so does the average WFA staining intensity per cell (Figure 3E). While gene expression analysis showed no change in most genes involved in PV cell activity (Figure 4F), CSPG biosynthesis (Figure 4G), PNN integrity (Figure 4H), and CSPG sulfation patterns (Figure 4I), we observed a decrease in *Pvalb* expression and a >6-fold increase in *Mmp9* expression, in keeping with some of the changes in *PV-scFvOtx2* mice at P400 (Figure 2H). The only other difference was increased expression of *Chst14* (Figure 3I), suggesting here again that the sequestration of OTX2 impacts the sulfation patterns of CSPGs. Taken together, these results suggest that blocking locally extracellular OTX2 indeed reverses the maturation level of PV cells in the PCx.

**Figure 3.**
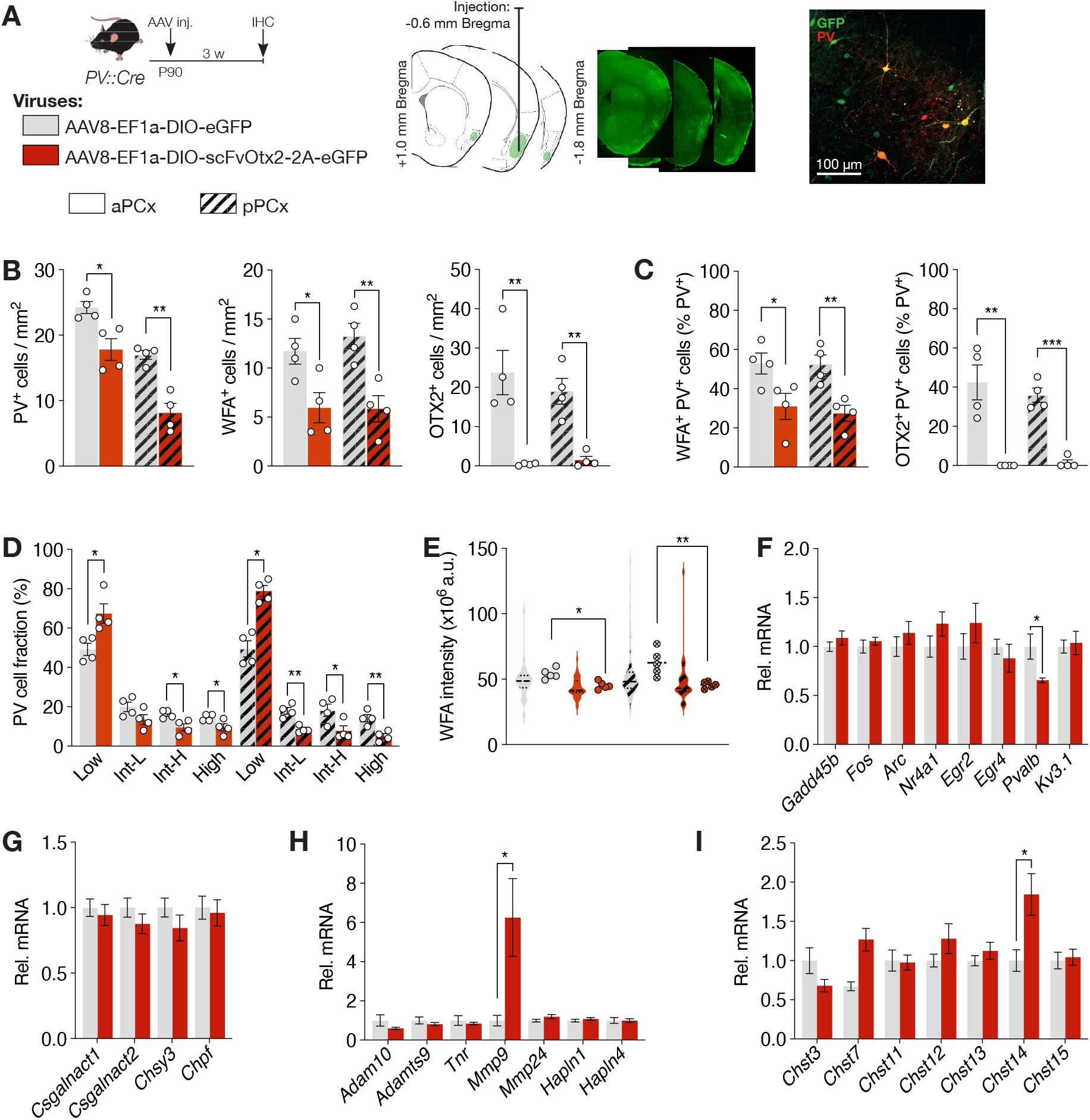
The piriform cortex undergoes remodeling after acute loss of function of OTX2. **(A)** The injection paradigm and virus design. **(B)** The number of PV+, WFA+, and OTX2+ cells in the PCx of P90 *PV::CRE* mice, 3-weeks after stereotaxic injection of virus expressing *GFP* (control) or *scFv-Otx2* (n = 4 mice per group). **(C)** The quantification of PV cell maturation by the number of PV+ cells stained for either WFA or OTX2 (n = 4 mice per group). **(D)** The distribution of PV staining level (n = 4 mice per group). **(E)** The staining intensity of WFA in PV+ cells in PCx (n = 5 mice per group). Violin plots show values for individual cells, while adjacent data points represent the average cell intensity per mouse. **(F-I)** Quantitative PCR analysis of genes for plasticity (F), CSPG biosynthesis (G), ECM components (H), and *Chst* genes (I) (n = 7-8 mice per group). All values ± SEM; *t*-test; \**P* < 0.05, \*\**P* < 0.01, \*\*\**P* < 0.001.

**Figure 4.**
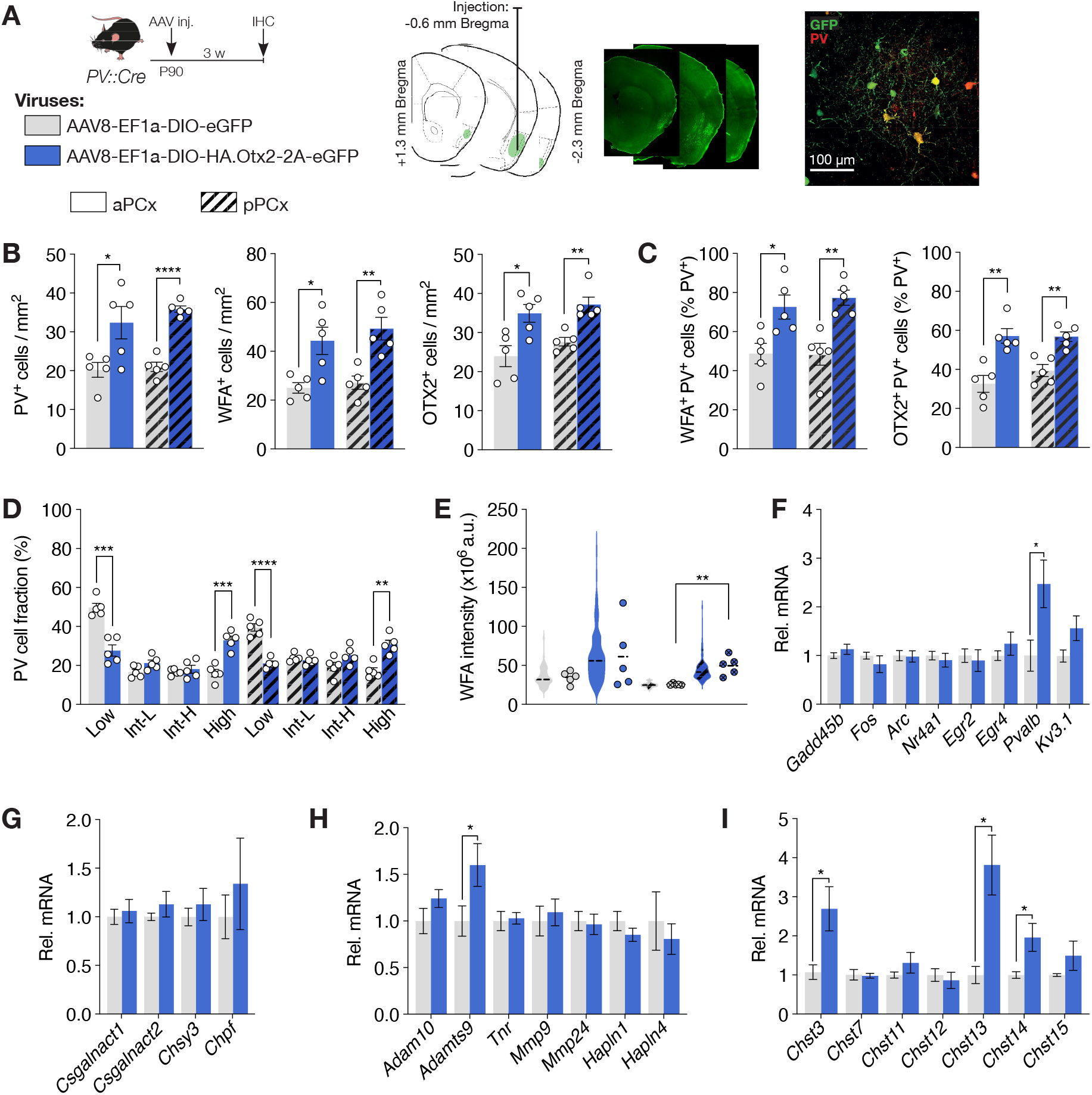
Acute OTX2 expression triggers PV cell maturation. **(A)** The injection paradigm and virus design. **(B)** The number of PV+, WFA+ and OTX2+ cells in PCx of *PV::CRE* mice, 3-weeks after stereotaxic injection of virus expressing *GFP* (control) or *Otx2* (n = 5 mice per group). **(C)** The quantification of PV cell maturation by the number of PV+ cells stained for either WFA or OTX2 (n = 5 mice per group). **(D)** The distribution of PV staining level (n = 5 mice per group). **(E)** The staining intensity of WFA in PV+ cells in PCx (n = 5 mice per group). Violin plots show values for individual cells, while adjacent data points represent the average cell intensity per mouse. **(F-I)** Quantitative PCR analysis of genes for plasticity (F), CSPG biosynthesis (G), ECM components (H), and *Chst* genes (I) (n = 6-12 mice per group). All values ± SEM; *t*-test; \**P* < 0.05, \*\**P* < 0.01, \*\*\**P* < 0.001.

### Acute OTX2 expression triggers PV cell maturation

To further investigate the role of OTX2 in PCx maturation, we used a conditional AAV8-EF1a-DIO-GFP-2A-Otx2 virus in *PV::Cre* mice for acute OTX2 over-expression. We observed a slightly wider distribution in GFP expression in PCx compared to loss-of-function injection in both aPCx (+1.34 ± 0.33 from bregma) and pPCx (−2.29 ± 0.16 from bregma) 3 weeks after P90 injection (Figure 4A). Interestingly, we found a pattern similar to OTX2 loss-of-function, albeit with an opposite sign. Forcing the local expression of OTX2 in PCx PV cells increases OTX2 levels as well as the number of PV+ cells and WFA+ cells, with significant increases of ~65% and ~80%, respectively (Figure 4B). This acute OTX2 expression triggers PV cell maturation, as shown by the increase in the number of PV+ cells surrounded by WFA or co-staining for OTX2 (Figure 4C), and by the significant shift in PV staining intensity towards the high-intensity population accompanied by a reduction of the low-intensity population (Figure 4D). WFA intensity increased in both regions, with significance in the pPCx (Figure 4E). Expression analysis of genes for PV activity (Figure 4F), CSPG biosynthesis (Figure 4G), PNN components (Figure 4H), and CSPG sulfation (Figure 4I) in PCx revealed an increase in *Pvalb, Adamts9* as well as *Chst3, Chst13*, and *Chst14*. These differences suggest OTX2 overexpression may alter PV activity and change PNN turnover and CSPG sulfation patterns.

### Impact of OTX2 in olfactory performance

To test whether acute changes in OTX2 levels within PV cells impacts PCx activity and function, we determined whether engaged olfaction by mice injected with either AAV8-EF1a-DIO-GFP-2A-scFvOtx2 or AAV8-EF1a-DIO-GFP-2A-Otx2 showed altered cFos staining compared to control mice at ~P110 (Figure 5A-D). Mice were sacrificed either at 1.5 h or 3 h after a 20-minute session involving random presentation of 4 different odors. While both cFos+ cell numbers and cFos staining intensity increased in the PCx compared to untreated control mice, the effect was significantly larger in pPCx (Figure 5B-C). We found that PV cells were activated by the stimulated olfaction. Approximately 20% of PV cells express cFos in unstimulated control conditions, while ~75% of PV cells in aPCx and pPCx express cFos 1.5 h and 3 h after odor presentation (Figure 5D), without affecting PV levels (Figure 5E). In mice injected with AAVs, we observed no change in cFos+ cell density in either aPCx or pPCx compared to controls at 3h, after the end of the session (Figure 5F). However, we observed a significant increase in cFos staining intensity in PV cells expressing scFvOTX2 (i.e., in GFP+ cells) in both aPCx or pPCx 3 h after odor presentation (Figure 5G), accompanied by a significant decrease in PV staining intensity within cFos+ cells (Figure 5H). Taken together, these results suggest that local OTX2 expression does not affect PV cell activity in the context of engaged olfaction of adult mice, while local disruption of OTX2 transfer, lowers PV levels and stimulates activity.

**Figure 5.**
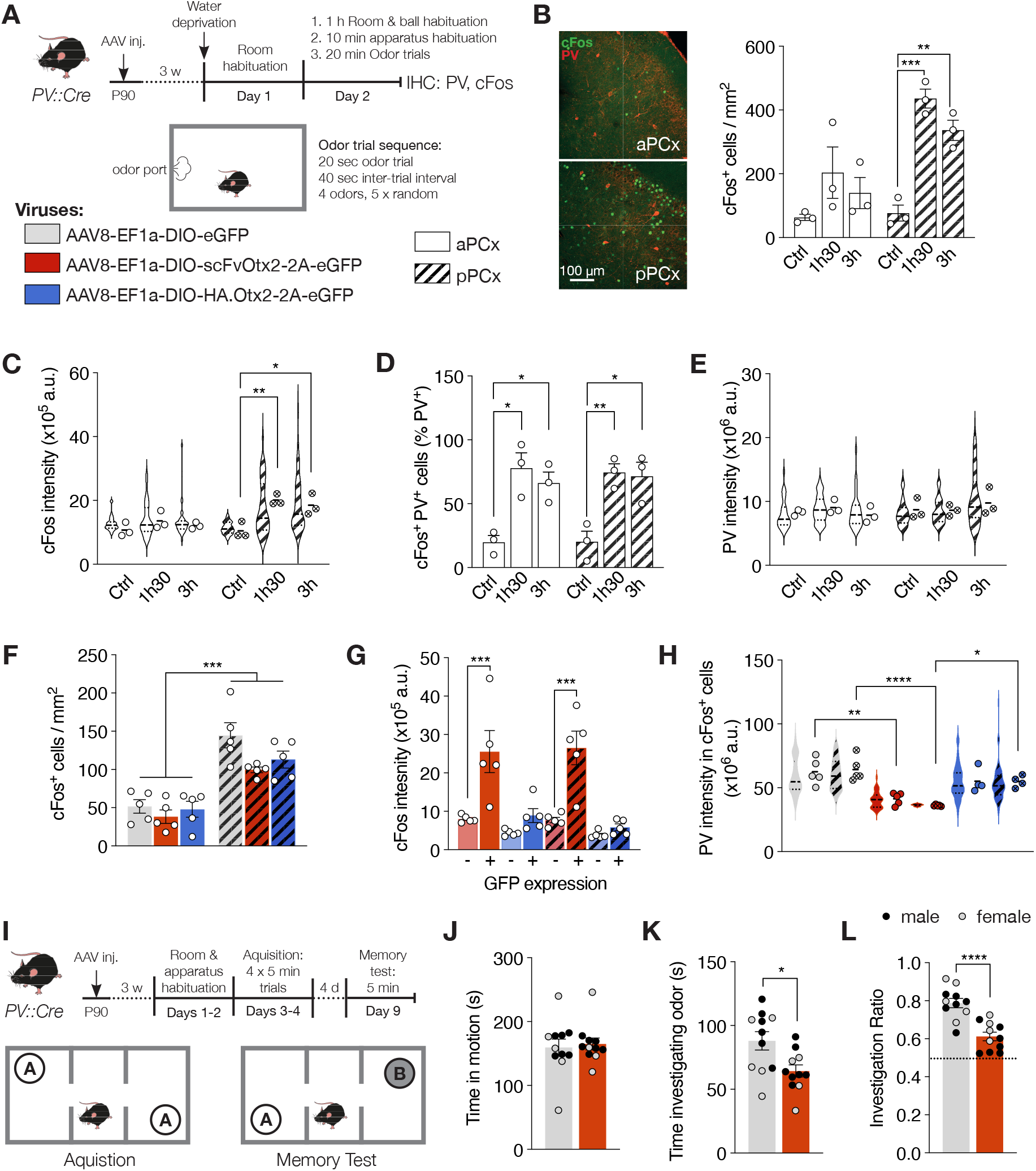
Altered olfactory memory through PV cell plasticity. **(A)** Experimental paradigm for odor representation. **(B)** Representative images of cFos and PV staining carried out in aPCx (white bars) and pPCx (striped bars) of untreated mice (control, CTL) and mice 1.5 or 3 h after odor presentation. Right panel: quantification of the number of cFos+ cells. **(C)** The percentage of PV+ cells co-stained with cFos. Violin plots show values for individual cells, while adjacent data points represent the average cell intensity per mouse. **(D)** The quantification of PV staining intensity. **(E)** The quantification of cFos staining intensity. **(F)** Quantification of the number of cFos+ cells in aPCx and pPCx, 3 h after odor representation in mice with virus expressing *GFP* (control), *scFvOtx2*, or *Otx2* (n = 5 mice per group). **(G)** cFos staining intensity in GFP- and GFP+ in aPCx and pPCx, 3 h after odor representation (n = 5 mice per group). **(H)** PV staining intensity in cFos+ cells in aPCx and pPCx, 3 h after odor representation (n = 5 mice per group). **(I)** Experimental paradigm for odor memory test. **(J)** Locomotor behavior during the memory test. **(K)** Time spent exploring both novel and familiar odorants during the memory test. **(L)** Investigation ratios representing time spent exploring the novel odor. Both mean ratios are significantly different from the chance level represented by the dashed line (n = 11 mice per group). All values ± SEM; one-way ANOVA, post hoc Tukey’s in (B-H); *t*-test in (J-L); \**P* < 0.05, \*\**P* < 0.01, \*\*\**P* < 0.001.

To test whether an acute decrease in OTX2 levels within PCx PV cells impacts animal behavior, we evaluated olfactory memory in mice injected with AAV8-EF1a-DIO-GFP-2A-scFvOtx2 compared to control mice. Olfactory performance was evaluated at ~P110 using a 3-chamber odor preference test (Figure 5I). Odor memory was acquired by 4 separate 5-minute sessions within 48 h during which the same odor was presented in both chambers. Odor memory was then tested 4 days later by changing the odor in one chamber and measuring the time spent near the odors in each chamber. While the loss-of-function mice showed similar locomotor activity compared to WT mice during the test (Figure 5J), they showed decreased overall olfactory interest by spending less time investigating the odors (Figure 5K). Importantly, they were significantly less interested in the novel odor compared to control mice (Figure 5L), suggesting that reducing OTX2 transfer to the PCx PV cells affects odor processing and decreases olfactory memory.

## Discussion

We find that PV cell trajectories in the PCx align more with associative areas than with neocortical primary sensory areas. In the P90 adult PCx, PV cell maturation (defined here by PV and PNN expression levels) and PV cell density are similar to higher order cortical areas, such as mPFC, but do not reach the elevated levels of V1. At P90, V1 is considered non-plastic while the mPFC is likely heterogeneous in its plasticity with multiple juvenile-like critical periods owing to its associative networks. Indeed, the non-patterned occurrence of PNNs suggests that PCx may experience multiple plasticity episodes, although these may be protracted given that significant maturation is only observed at P400. Furthermore, our analysis of the *PV-tdTom* mouse model revealed heterogeneity in the PCx PV cell population, as PV cells expressing TOM had significantly higher levels of PV and PNNs. This heterogeneity may be developmental in origin, as suggested by the segregation in TOM expression, may arise from delayed differentiation, a possibility given the existence of “immature” neurons in PCx (Ghibaudi et al., 2023), and/or may be due to different PCx microcircuits (Jiang et al., 2021). These local circuits would depend on the nature and source of long-range inputs, such as olfactory area nucleus, BLA, mPFC, HC and/or EC (Ryu et al., 2021), although we did not discern a layer-specific pattern of PV cell activation within either aPCx or pPCx. PV cells are also sensitive to neuromodulators, with dopamine acting on D1-class dopamine receptors on PV cells in the PCx to increase their intrinsic excitability (Potts and Bekkers, 2022). Future tracing studies may shed light on whether distributed yet specific microcircuits involving PV cells are stabilized and matured by specific inputs from olfactory areas, PCx or other brain regions.

The reduced PV cell maturation in the PCx of P400 *PV-scFvOtx2* mice and the direct influence of altered OTX2 levels on PV and PNN staining point to a role for OTX2 in regulating PCx PV cell maturation. Such OTX2 activity is complex and can occur at several levels, including signaling, translation, transcription, and epigenetics (Apulei et al., 2019; Hou et al., 2017; Lee et al., 2017; Sakai et al., 2017). Long-term reduction of OTX2 in the PCx results in increased expression of *Adamts9* and *Mmp9*, which proteolyze proteoglycans (Huntley, 2012; Lemarchant et al., 2013) and would account for lower PNN levels through matrix reorganization in P400 mice. We also observed increased expression of *Chst7, Chst11*, and *Chst13*, which may alter the pattern of CSPG sulfation (Mikami and Kitagawa, 2013). In the acute OTX2 loss-of-function AAV-scFvOtx2 mice, we find increased expression of *Mmp9* and *Chst14*, while in the acute OTX2 gain-of-function mice, we find increase expression of *Adamts9, Chst3, Chst13* and *Chst14*. There is a known feedback loop between OTX2 accumulation and PNN formation (Di Nardo et al., 2018), and our findings further suggest that OTX2 may also affect CSPG sulfation patterns. Overall, these changes in gene expression may account for the observed changes in WFA staining in the different models, and confirm that MMP9 plays a role in reducing PNN levels (Kelly et al., 2015; Ribot et al., 2021) while ADAMTS9 allows for greater PNN accumulation, in keeping with its expression during V1 critical period and suggested role in allowing for PNN remodeling and higher density (Apulei et al., 2019). However, these genes are not PV cell specific and expression levels do not correlate with OTX2 levels, strongly suggesting that OTX2 does not directly regulate their expression but instead regulates upstream transcription factors and affects signaling to neighboring neurons and glial cells. Future DNA and RNA sequencing studies focused on PCx PV cells will be required to more robustly assess the direct impact of OTX2 on epigenetics, gene expression and active translation.

Our experiments are in keeping with prolonged plasticity and age-related maturation of PCx neurons and circuits (Ghibaudi et al., 2023; Wu et al., 2020). By directly expressing *Otx2* in PCx PV cells, not only did *Pvalb* expression, PV staining and PNN accumulation increase, but also the number of PV+ cells, suggesting that some PV cells are in a form of quiescence but nevertheless available when solicited. Furthermore, changes in PV cell maturation were found to be region-dependent. Although viral expression was uniform throughout PCx, the effect of both loss- and gain-of-function paradigms was strongest in the pPCx. A confounding issue may be the use of the *PV::Cre* mouse model for conditional expression of scFvOTX2 and OTX2 in PV cells, given that CRE is expressed in only ~50% of PCx PV cells, which also tend to be the more “mature”, as revealed by the *PV-tdTom* mouse. This heterogeneity may not be ideal for our gain-of-function experiments, as more “mature” cells will be targeted, even though we cannot discount local OTX2 transfer between PV cells. While we observed increased PV and PNN levels, we did not observe changes in cFos activity. This may not have been the case had the pool of less mature cells been targeted. However, for our loss-of-function experiments, this issue is less relevant, as scFvOTX2 is secreted and will sequester extracellular OTX2 and affect neighboring PV cells throughout the PCx. Indeed, the conditional expression of scFvOTX2 led to changes in PV and PNN levels, cFos activity, and behavioral outcomes.

OTX2 levels in PCx PV cells are likely affecting plasticity states that impact olfactory-dependent behavior in adult mice. Our behavioral paradigm consisted in several acquisition sessions followed by a memory test 4 days later. It is therefore possible that the poorer performance of mice with reduced OTX2 levels is related either to a learning deficit during the acquisition sessions or to a memory defect. At this juncture, we do not know whether or not there are discrete critical periods of plasticity in the PCx. We have previously proposed a two-threshold model of critical period plasticity timing based on the level of OTX2 accumulation in PV cells (Prochiantz and Di Nardo, 2015; Spatazza et al., 2013), and given the sensitivity of PV cells to OTX2 levels, it is likely that PCx plasticity is also based on similar molecular pathways but that the kinetics of OTX2 accumulation are slower. Lower levels of OTX2 induced by acute scFvOTX2 expression may keep PCx PV cells in a highly plastic state, negatively impacting odor memory acquisition and/or retention.

How PV staining levels correlate with plasticity and affect behavior remains unknown, but a recent study using a mouse model for PV-cell-specific deletion of *Ntrk2* (gene for TrkB protein) showed delayed PV cell maturation and altered cortical networks in the PCx (Lau et al., 2022). While the impaired PV cell function resulted in aberrant spike patterns of PCx pyramidal cells in response to sensory input and in a paradoxical decrease in overall network excitability (Lau et al., 2022), it remains possible that this impairment may be due to global changes in brain function given that the PV cell deletion of TrkB is brain-wide in their model. Another recent study involving optogenetic stimulation of slice preparation showed that recurrent inhibition from aPCx PV cells gated output in a layer-specific manner and that functional plasticity driven by enhanced PV cell inhibition of superficial pyramidal cells (Jiang et al., 2021). Indeed, inactivation of aPCx PV cells can increased pyramidal cell firing rates and promote long-term potentiation (Canto-Bustos et al., 2022). Thus, PV cells have the potential to continually shape PCx output in response to sensory input. However, how this potential may change during aging remains to be addressed. Morphological changes in PV cells could alter the extent of feedback connections while PNNs may restrict functional plasticity and favor sub-networks with long-range connections to other olfactory or associative brain regions.

Further work is needed to determine the role of PV cell activity for odor-driven behavior. Hints may come from other associative brain regions, such as mPFC or HC, where direct inhibition of PV cell activity can activate circuits that alter fear behavior (Courtin et al., 2014; Donato et al., 2013). In the aPCx, PV cell activity is modulated by disinhibitory circuitry driven by VIP interneurons (Canto-Bustos et al., 2022), while similar VIP-controlled circuits exist in the HC for fear behavior and learning (Donato et al., 2013) and in the primary motor and visual cortices during motor learning (Donato et al., 2013; Fu et al., 2014). However, these forms of learning and plasticity may not be relatable to critical period plasticity given that VIP-mediated disinhibition is not implicated in critical period timing in sensory cortices (Takesian et al., 2018). Previous studies have shown rostro-caudal asymmetry in inhibition levels within aPCx, with a prominent role for somatostatin interneurons (Large et al., 2018; Luna and Pettit, 2010). While we do not find differences in PV cell density or staining levels across the aPCx, nor between the aPCx and pPCx, the pPCx seems to be more plastic, with higher cFos levels after odor stimulation that are amplified in PV cells upon reduction of OTX2 levels. Taken together, our findings show that OTX2 activity in the PCx is indeed affecting PV cell maturation states and plasticity impacting odor behavior, but that the PCx PV cell population is much more heterogeneous compared to sensory cortex PV cell populations and likely not subjected to strictly defined windows of plasticity.

## Acknowledgements

We would like to thank Philippe Mailly for essential help with image analysis, Yves Dupras for constructing the behavior apparatus, Clémentine Pesnel for technical help, David Benacom for scientific discussions, and Alain Prochiantz for scientific discussions and critical reading of the manuscript. Funding was provided by the Fondation Bettencourt Schueller and the Agence Nationale de la Recherche (ANR-18-CE16-0013-01).

## Materials & Methods

### Animal management

All animal housing and experimental procedures were carried out in accordance with the recommendations of the European Economic Community (2010/63/UE) and the French National Committee (2013/118). This research (project n° 19086) was validated by the regional ethics committee (CEA 59) and authorized by the French Ministry of Higher Education, Research and Innovation. Adult C57BL/6J wildtype (WT) mice were obtained from Charles River Laboratories, while *PV::Cre* (strain n° 017320) and *Rosa-tdTomato* (*Ai14* strain n° 007914) reporter mice were generated from JAX founders. The *scFvOtx2*^*tg/o*^ mice were generated as previously described (Bernard et al., 2016). The double transgenic crosses (*PV::Cre;Rosa-dtTomato* and *PV::Cre;scFvOtx2*^*tg/o*^) were generated in-house. Mice were housed with *ad libitum* access to food and water and under a 12h light/dark cycle. For surgical procedures, animals were anesthetized with xylazine (Rompun 2%, 5 mg/kg) and ketamine (Imalgene 500, 80 mg/kg) by intraperitoneal injection.

### Stereotaxic injections

Adeno-associated viruses (AAV) were produced by Vector Biolabs (Malvern, PA, USA): AAV8(EF1a)-DIO-scFvOtx2-2A-GFP and AAV8(EF1a)-DIO-GFP-2A-Otx2. Unilateral or bilateral stereotaxic injections (bregma: × = 3.8 mm, y = ± 0.6 mm, z = 4 mm) of 0.5 μl high-titer AAV (~10^13^ GU/ml) were performed at a rate of 0.1 μl/min with a Hamilton syringe. Mice were used for histological and behavioral analysis at least 3 weeks after infection.

### Quantitative RT-PCR

Mice were sacrificed by cervical dislocation and PCx was micro-dissected in ice-cold phosphate buffered saline (PBS) and frozen on dry ice. Total RNA was extracted will AllPrep DNA/RNA Mini Kit (Qiagen). cDNA was synthesized from total RNA with QuantiTect Reverse Transcription kit (Qiagen). Quantitative PCR reactions were carried out in triplicate with SYBR Green I Master Mix (Roche) on a LightCycler 480 system (Roche). Expression was calculated by using the 2^-ΔΔCt^ method with *Gapdh* as housekeeping reference gene. Primer sequences:

*Adam10*, F-GGGCTCTCCATGTAATGACTTC, R-CACAATCCACTCAGCAATGTTT

*Adamts9*, F-GTAAGCACCTTCCTAAGCCAC, R-GATCTGACAGCCCACATGCCTC

*Tnr*, F-TTGTGAGAGACAGCAGAGCG, R-TCTGTCTCTGCGTGTTGAGC

*Mmp9*, F-CATAGAGGAAGCCCATTACAGG, R-CCTGTCTACACCCACATTTGAC

*Hapln1*, F-ATGTAAGCGGAAGGCATTCTAA, R-AATCCTGGTGATTCTCAGCCTA

*Hapln4*, F-GGTGCACTGCAGACT, R-AGCGACCAAGAACCACAACA

*Chst3*, F-CCCCAAGATGGTCTCGTGTT, R-GCCCAGGCCCTGATTTTAGT

*Chst7*, F-GGGACTCGTCGAGGACAAAG, R-CCTTTAGGTTGAGGCCTGGG

*Chst11*, F-AAAGTATGTTGCACCCAGTCAT, R-GGATGGGATTGTAGAGCTCCTG

*Chst13*, F-TTTTCCAGGACATCAGCCCC, R-ACACCCTTATTGCAGTCGCA

*Chst14*, F-CATCCTATCGGAGATGAAACCC, R-GATAGGCCTCTAGGCTTAGGAT

*Chst15*, F-CATTCCTGACCCAAGACTTCAT, R-AGTCTGAGTACAACCTCTCCAC

*Gapdh*, F-TGACGTGCCGCCTGGAGAAAC, R-CCGGCATCGAAGGTGGAAGAG

### RNAscope fluorescent in situ hybridization

Mice were given a lethal dose of Euthasol and perfused transcardially (5 ml/min) with 15 ml of PBS followed by 15 ml of 4% paraformaldehyde (PFA) in PBS. Dissected brains were post-fixed in 4% PFA, PBS at 4 °C overnight and then rinsed extensively in PBS. Coronal vibratome sections (40 μm) were incubated with RNAscope hydrogen peroxide solution from Advanced Cell Diagnostics (ACD) for 10 min at room temperature (RT), rinsed in Tris-buffered saline with Tween (50 mM Tris-Cl, pH 7.6; 150 mM NaCl, 0.1% Tween 20) at RT, collected on Super Frost plus microscope slides (Thermo Scientific), dried at RT for 1 h, rapidly immersed in ultrapure water, dried again at RT for 1 h, heated for 1 h at 60°C and dried at RT overnight. The next day, sections were immersed in ultrapure water, rapidly dehydrated with 100% ethanol, incubated at 100 °C for 15 minutes in RNAscope 1X Target Retrieval Reagent (ACD), washed in ultrapure water and dehydrated with 100% ethanol for 3 minutes at RT. Protease treatment was carried out using RNAscope Protease Plus solution (ACD) for 30 minutes at 40°C in a HybEZ oven (ACD). Sections were then washed in PBS before *in situ* hybridization using the RNAscope Multiplex Fluorescent V2 Assay (ACD). The PV probe (Mm-*Pvalb*-C2, ACD catalog #421931-C2) was hybridized for 2 h at 40 °C in a HybEZ oven, followed by incubation with signal amplification reagents according to manufacturer instructions. The images were acquired with a W1-CSU spinning-disk ZIESS Axio Observer Z1 microscope.

### Immunohistochemistry

Anesthetized mice were perfused transcardially (5 ml/min) with 15 ml of PBS and 15 ml of 4% PFA, PBS. Dissected brains were post-fixed in 4% PFA, PBS at 4 °C overnight and then rinsed extensively in PBS. Fluorescent immunohistochemistry was performed on coronal vibratome sections (40 μm). Briefly, after permeabilization and blocking with PBS, 1% Triton, 5% fetal bovine serum (FBS) for 45 min at RT, sections were incubated in PBS, 1% Triton, 5% FBS overnight at 4 °C with primary antibodies: anti-OTX2 (mouse monoclonal, in house, 1/50); anti-PV (rabbit polyclonal, Swant, 1/500); biotinylated *Wisteria floribunda* agglutinin (WFA; Sigma L1516, 1/500); anti-GFP (chicken polyclonal, Abcam, 1/800); and anti-cFos (rabbit monoclonal, Cell Signaling, 1/500). After 3 washes in PBS, secondary antibodies (Alexa Fluor-conjugated, Molecular Probes, 1/2000) were incubated for 90 min at RT. After 3 washes in PBS, sections were mounted in DAPI-fluoromount medium and kept at 4 °C. Images were acquired with either a ZEISS Axio Zoom.V16, Leica DM16000 SP5 confocal, or W1-CSU spinning-disk ZIESS Axio Observer Z1 microscope. Cell number and staining intensity were quantified with ImageJ and in-house macros.

### Behavior analyses

#### Odor memory test

The 3-chambered testing apparatus consisted of a grey-colored PVC arena (dimensions *l* × *d* × *h* in cm: 78 × 45 × 30) with a smaller central chamber and two external chambers. Two cylindrical cages (10.2 cm in diameter and 18 cm tall) containing odor balls were placed in the external chambers at least 10 cm away from the walls. The cages and the arena were thoroughly cleaned with 70 % ethanol between each trial. Clean 5 cm wooden balls containing a cotton swab were introduced into the cylindrical cages just prior to acquisition. The odors (Sigma Aldrich), 0.3% ethyl-butyrate and 0.3% limonene, were diluted in mineral oil (Sigma Aldrich). For habituation, the mice were kept in their home cage and spent 4 h on Day 1 in the testing room without the experimenter. On Day 2, the mice spent 4 h in the testing room with the experimenter and were placed individually for 10 min in the arena without odor cages. Mice were returned to the housing area after each habituation phase. During the acquisition phase on Days 3 and 4, the mice were individually placed in the center chamber with both cages containing the same odor. Mouse movement was recorded for 5 min in the absence of the experimenter. On Days 3 and 4, two 5-min acquisition trials were performed spaced 3 h apart to provide a total of four acquisition trials. Mice were returned to the housing area after the daily trials.

For the memory test, the mice were water-restricted on Day 8 in order to increase motivation. On Day 9, mouse movement was recorded for 5 min in the arena with one chamber containing the acquisition odor and the other chamber containing a novel odor. The cages and the arena were thoroughly cleaned with 70% ethanol between each trial, and the location of the novel odor was alternated between chambers for subsequent trials. Odor investigation time was defined as the amount of time the snout was directed towards an odor cage (within 45°) within a 2 cm radius. Novel odor preference was defined as the ratio of novel-odor to total-odor investigation time.

#### Odor-induced protein c-Fos staining

Odors were delivered using a 4-channel olfactometer (Automate Scientific). Clean air (pressure, 3 atm) was directed to 4 individual air lines each connected to a 10 mL glass bottle containing mineral oil (Sigma Aldrich) in which the odors were diluted. The bottles themselves were connected via Tygon tubing to computer-controlled (Arduino) solenoid valves (Valvelink, Automate Scientific). Four different freshly-prepared odors (Sigma Aldrich) were tested: empty bottle, 0.3% ethyl-butyrate, 0.3% hexanal, and 0.3% limonene. The testing apparatus consisted of a clear polycarbonate box (dimensions *l* × *d* × *h* in cm: 37.5 × 23 × 10.5) with a 5 cm wooden ball placed at the exit of a scent port controlled by the olfactometer. The box was thoroughly cleaned, inside and out, with 70 % ethanol at the end of each trial. Clean wooden balls were introduced into the box at the beginning of each acquisition.

The mice were habituated to the testing room and the experimenter 24 h before the day of testing and were water-deprived to increase motivation. On the day of the trial, the mice were kept in their home cage and habituated to the wooden balls in the testing room. During the pre-trial phase, the mice were individually placed in the box for 10 min. They were then presented with each of the 4 odors 5 times randomly for 20 sec (trial) spaced by 40 sec (inter-trial interval, ITI). Mice were sacrificed by perfusion at either 0, 1.5, or 3 h after the end of the trial phase for histological analysis of cFos staining.

### Statistical Analysis

All statistics were performed using Prism (version 9.3.0, GraphPad Software). Main effects and interactions with more than 2 groups were determined using analyses of variance (ANOVA) or 2-way ANOVA with Tukey’s multiple comparison *post hoc* test. The *t*-test was used for single comparisons among two groups.

